# Classes of ITD predict outcomes in patients with AML treated with *FLT3* inhibitors

**DOI:** 10.1101/322354

**Authors:** Gregory W. Schwartz, Bryan Manning, Yeqiao Zhou, Priya Velu, Ashkan Bigdeli, Rachel Astles, Anne W. Lehman, Jennifer J.D. Morrissette, Alexander E. Perl, Martin Carroll, Robert B. Faryabi

## Abstract

Recurrent internal tandem duplication (ITD) mutations are observed in various cancers including acute myeloid leukemia (AML). ITD mutations of Fms-like tyrosine kinase 3 (*FLT3*) receptor increase kinase activity, and are associated with poor prognostic outcomes. Currently, several small-molecule *FLT3* inhibitors (FLT3i) are in clinical trials for targeted therapy of high-risk FLT3-ITD-positive AML. However, the variability of survival following FLT3i treatment suggests that the mere presence of *FLT3*-ITD mutations in a patient might not guarantee effective clinical response to targeted inhibition of *FLT3* kinase. Motivated by the heterogeneity of *FLT3*-ITD mutations, we sought to investigate the effects of *FLT3*-ITD structural features on response to treatment in AML patients. To this end, we developed HeatITup (HEAT diffusion for Internal Tandem dUPlication), an algorithm to efficiently and accurately identify ITDs and classify them based on their nucleotide composition into newly defined categories of “typical” or “atypical”. Typical ITDs insert sequences are entirely endogenous to the *FLT3* locus whereas atypical ITDs contain nucleotides exogenous to the wildtype *FLT3*. We applied HeatITup to our cohort of *de novo* and relapsed AML patients. Individuals with AML carrying typical ITDs benefited significantly more from FLT3i than patients with atypical ITDs, regardless of whether FLT3i was used after initial induction or at relapse. Furthermore, analysis of the TCGA AML cohort demonstrated improved survival for patients with typical ITDs treated with induction chemotherapy. These results underscore the importance of structural discernment of complex somatic mutations such as ITDs in progressing towards personalized treatment for AML patients.

## Introduction

Several recurrent somatic genetic alterations in acute myeloid leukemia (AML) have been incorporated in the standard-of-care recommendations for diagnosis, response assessment, and treatment outcome prediction (1,2). Internal tandem duplication (ITD) mutations in the Fms-like tyrosine kinase 3 (*FLT3*) gene are among the most frequent somatic alterations in AML (3). *FLT3*-ITDs are always in-frame insertions and result in the loss of auto-inhibitory function, leading to increased tyrosine kinase activity. The presence of *FLT3*-ITD, regardless of the cytogenetic classification, is associated with significantly increased relapse risk and decreased overall survival (4–12). Hence, screening for the presence of the *FLT3*-ITD mutation is recommended (4,7,8) and genetic tests to identify *FLT3*-ITD mutations are now part of routine diagnostic workups for both *de novo* and relapsed AML patients. Recently, next-generation sequencing (NGS) diagnostic assays have been developed to test for the presence of mutant *FLT3*-ITD in AML patients (13,14). Although these new technologies enable a more detailed analysis concerning the effect of complex mutation structures on clinical outcome, we still lack an understanding of the heterogeneity of ITD structures and its impact on the clinical outcomes. We present the first study to collate the effect of structural properties of *FLT3*-ITDs beyond the length of the duplicated segment itself on response to targeted therapy.

The prevalence of prognostically high-risk *FLT3*-ITD mutations resulted in the introduction of *FLT3* inhibitory agents as therapeutic options for *FLT3*-ITD-positive AML. More recently, *FLT3* tyrosine kinase inhibitory small-molecules (FLT3i) with various levels of specificity and potency such as Sorafinib (15,16), Quizartinib (AC220) (17–20), Midostaurin (21,22), and Gilteritinib (ASP2215) (23,24) have been developed. Although the newer generation of FLT3i such as Quizartinib provides stronger and more prolonged anti-leukemic activity, the complete response rate still is at most 50% (depending on response criteria) (20–28), demonstrating that much remains to be learned before effective use of these agents for targeted therapy in *FLT3*-ITD-positive patients. One possibility for this variability in treatment response may be due to the differences in the structure of ITD mutations.

Unlike many other hotspot mutations, ITD mutations are diverse and complex genetic alterations (29,30). While in-frame ITDs are mainly in exon 14 of *FLT3*, they exhibit marked structural heterogeneity. For example, diversity in the ITDs duplicated sequence length, position of duplications, or mutation burden are reported (31). Earlier studies using fluorescence-based capillary gel electrophoresis (CGE) suggested that the ITD length could impact the prognostic outcome of AML patients treated with chemotherapy-based regimens (11,29). However, the link between the response to FLT3i and *FLT3*-ITD structural features remains unknown. We hypothesized that the *FLT3*-ITD structural features beyond the length impacts clinical response to FLT3i treatment in *de novo* and relapsed AML patients. A prerequisite to test this hypothesis was an analytical method to characterize the complexity of ITD somatic mutations from NGS readouts.

To address the unmet need for accurate identification of ITD mutations from the next-generation sequencing (NGS) readouts and further characterize their complexity, we developed HeatITup (HEAT diffusion for Internal Tandem dUPlication). HeatITup is an efficient and easy to use open source software that is available for download from https://github.com/faryabib/HeatITup. We retrospectively applied HeatITup to samples from both the Cancer Genome Atlas (TCGA) cohort and AML patients sequenced at the University of Pennsylvania (UPENN) to identify and annotate *FLT3*-ITD mutations. Based on the divergence in the nucleotide composition of ITD mutations beyond the duplicated segment, we introduced two distinct classes of “typical” and “atypical” ITD mutations. An ITD mutation was classified as typical if the insertion is entirely consisting of nucleotides endogenous to wild-type *FLT3* locus, otherwise the ITD was classified as atypical.

Equipped with our refined ITD classification, we systematically studied the impact of the presence of *FLT3*-ITD mutations and their structural features on the clinical response to *FLT3* inhibitory agents in *de novo* and relapsed AML patients. Our data showed that AML patients carrying typical ITD mutations with insert sequences entirely endogenous to the *FLT3* locus benefit significantly more from FLT3i than patients with atypical ITDs whose mutations consist of nucleotides mainly exogenous to the wild-type *FLT3*. Our observation demonstrates that the selectivity of response to FLT3i depends not only on the presence of an ITD mutation in the AML genome but is also a function of the ITD structure beyond the mutation length.

## Materials and Methods

### The University of Pennsylvania (UPENN) cohort

318 amplicon-based NGS clinical samples from 216 AML patients were analyzed. These samples were collected from 2013 to 2017. Data collection was frozen in January 2017. 64 patients were sequenced more than once throughout the course of their disease. Overall, 113 *FLT3*-ITD-positive samples from 71 individuals with retrievable and reliable clinical outcome annotations in electronic health record (EHR) systems of the University of Pennsylvania were identified. Structural characteristics of FLT3 mutations were characterized in these 71 patients. 42 samples were cytogenetically characterized at the same time as *FLT3* analysis, among them 2.4% were favorable, 81.0% were intermediate, and 16.7% were unfavorable. 58 out of 71 patients who received a FLT3 inhibitor (FLT3i) as part of their therapy were selected and used to assess the correlation between *FLT3* mutation type and survival after therapy (Table 1). Stringent conditions were used to further annotate this cohort to minimize clinical variability across the individuals included in the survival analysis. Accordingly, the patients were stratified as either *de novo* or relapsed AML. The *de novo* group consisted of individuals diagnosed with *de novo* AML who received induction chemotherapy and the NGS-based diagnostic test within 30 days of the diagnosis date (n = 32). Alternatively, the relapsed group consisted of individuals with relapsed AML who received induction chemotherapy prior to relapse with an observable *FLT3*-ITD mutation within 30 days of the first relapse diagnosis date (n = 26).

### DNA sequencing

The protocol for this study was approved by the University of Pennsylvania Medical School Institutional Review Board. *FLT3* sequencing was performed via targeted next-generation sequencing (HUP-HemeV2; the University of Pennsylvania, Philadelphia, PA) on the MiSeq sequencing system (Illumina) as described before (13). HUPHemeV2 is a CLIA-certified NGS assay designed at the University of Pennsylvania to detect somatic mutations in *FLT3* (13). DNA was quantified using a fluorescent based measurement (Qubit, Life Technologies) and 20 to 250 ng of DNA was used for enrichment. Following library preparation with the TruSeq Amplicon assay (IIlumina), libraries were pooled and sequenced on the MiSeq system to an average depth of coverage greater than 1,000x.

### The Cancer Genome Atlas (TCGA) AML cohort

Sequencing files for 321 samples from 149 individuals with AML in the TCGA cohort were downloaded from the Genome Data Commons (https://gdc.cancer.gov) (3) and analyzed.

### ITD Clonal assignment

All reads with a similar *FLT3*-ITD mutation were assigned to a clone with an allele frequency (AF). We provided a helper program to assist in grouping together similar mutations into clones. We removed clones with an AF smaller than 1%, resulting in 135 typical and 39 atypical clones in 71 individuals. Due to lower sequencing coverage of the TCGA samples, we only included clones if their *FLT3*-ITD duplication length was 15 or greater. Each individual’s representative *FLT3*-ITD clone was determined based on the dominant clone from the earliest sample in the UPENN cohort or, due to less coverage, the longest duplication in the TCGA cohort. See Supplementary Information for further information.

### Statistical Analysis

The Mann-Whitney U test was used in comparing duplicated and spacer segment lengths, as well as purine and pyrimidine abundance. Exogenous nucleotide abundance was compared using Tukey’s HSD after ANOVA. For 58 patients with *de novo* or relapsed AML, the date of diagnosis and the date of censor (death or last recorded follow-up) were obtained from the University of Pennsylvania EHR and verified using Leukemia Tissue bank database of the University of Pennsylvania. Overall survival (OS) was determined using Kaplan-Meier survival curves. Survival statistical analyses were performed using stratified Cox regression (32) to control for disease stage (*de novo* and relapsed) with sex, age, allogenic transplant, remission at initiation of FLT3i treatment, and FLT3i type (Sorafenib:30, Gilteritinib:32, Crenolanib:1, Quizartinib:9, PLX3397:9, FLX925:1, Midostaurin:1) as additional covariates. 21 patients received more than one inhibitor. FLT3i type was categorized into four major groups: Sorafenib, Gilteritinib, both Sorafenib and Gilteritinib, or Other for statistical analysis. The proportional hazards assumption for each variable was tested and met using Schoenfeld residuals (33).

## Results

Unlike many other hotspot mutations, ITDs are diverse and complex genetic alterations. We observed that the inserted segment of DNA could appear immediately next to its origin of duplication or there could be intervening nucleotides (Figures 1a and 3). While ITD mutations are mainly in the exon 14 of *FLT3*, the insertion location, the length and nucleotides of the duplicate segment, and the composition of the DNA segment between the duplication and its origin add to the ITD structural complexity (Figures 1a and 3).

We defined several attributes that capture ITD diversity. We termed the part of an inserted DNA segment that is a duplication of the reference as “duplication”, and the nucleotides in-between the duplication and its origin as “spacer” (Figure 1a). We observed that a spacer could consist of nucleotides that were either endogenous to the *FLT3* wild-type locus or were not part of the wild-type exon (Figures 1a, 3). We classified an ITD as “atypical” if the spacer sequence consisted of at least one nucleotide exogenous to the *FLT3* wild-type sequences, otherwise the ITD was classified as “typical” (Figures 1a, 3). Importantly, while atypical and typical ITDs could have spacer sequences, they both resulted in in-frame mutations.

Since ITD mutations are a combination of alterations, commonly used tools such as Pindel (34) do not directly detect them. To address the unmet need for accurate identification of ITD mutations from NGS short sequence reads, and further characterize the complexity of ITD mutations based on the aforementioned structural attributes, we developed and applied HeatITup to *FLT3* gene sequences in AML patients. Rather than search for an insertion like methods proposed in (3,34–36), HeatITup analyzes both mapped and unmapped short-read sequences and detects the largest duplicated sequences; characterizes the sequence composition of the spacer segment; and classifies ITDs as typical or atypical (Figures 1a and 3, see Supplemental Information). By repeated usage of suffix trees with “end mutations” (see Methods and Materials, and Supplemental Information), HeatITup is able to detect both the duplications as well as any possible point mutations within them (Figures 1b and c). HeatITup also analyzes the composition of spacer sequences by using a combination of a Hamming distance comparison with a reference sequence and discrete Gaussian kernel-based heat diffusion and detects the sequence segment within the spacer exogenous to the wild-type locus (Figures. 1b and c, Methods and Materials, and Supplemental Information). As such, HeatITup is the first method capable of annotating duplication, spacer, and exogenous segments in addition to detecting ITDs.

To assess the accuracy of HeatITup, we performed a comparative analysis against four commonly used methods based on 321 samples from 149 individuals profiled in the TCGA AML cohort (3). These samples were previously scored for the presence of ITD mutations in *FLT3* by the method used in the original analysis of the TCGA AML cohort (3), ITD Assembler (37), Pindel (34), and Genomen (38). HeatITup found 7 new *FLT3*-ITD-positive individuals missed by these methods, exhibiting that HeatITup accurately detects ITD mutations while providing higher sensitivity than these methods (Figure 2). We identified a total of 35 *FLT3*-ITD-positive individuals, where 20 harbored typical and the remaining carried atypical ITDs. Compared to the earlier analysis, HeatITup increased the percentage of detected *FLT3*-ITD-positive cases in the TCGA cohort from 19% to 23%, which is closer to the expected frequency of *FLT3*-ITD-positive cases in a representative AML cohort based on other methodologies such as CGE.

We then analyzed amplicon-based NGS of 216 patients in the UPENN cohort (see Material and Methods). HeatITup identified 113 *FLT3*-ITD-positive samples from 71 individuals (Figure 2). Our analysis improved the detection of *FLT3*-ITD and found an additional 5 *FLT3*-ITD-positive individuals that were reported as *FLT3*-WT by the analysis previously performed using GATK/Pindel-based Clinical Laboratory Improvement Amendment (CLIA)-confirmed analysis workflow.

We next assessed the accuracy of HeatITup to classify samples as typical or atypical. We analyzed 32 UPENN samples with HeatITup and compared the results with experienced genetic reviewers who manually verified the presence and classified ITD mutations. 38 of 39 ITDs were classified concordantly while one sample, CPDC140113, had a discordant classification (Figure 3, c.f. red-boxed individual). Upon further inspection, we found that the misclassified *FLT3*-ITD in sample CPDC140113 was indeed atypical and HeatITup was able to detect and correct the error of manual classification. Together, these evaluations exhibited the precision of HeatITup in detecting and classifying ITD mutations accurately based on their nucleotide composition into typical or atypical categories while providing high sensitivity.

We quantified the clonality and structural characteristics 113 *FLT3*-ITD-positive samples from 71 individuals in the UPENN cohort to assess the structural heterogeneity of ITD mutations (see Material and Methods). We found that our cohort consisted of similar proportions of mono-and polyclonal cases (Figure 4a). While 54% of individuals harbored one *FLT3*-ITD clone, 4% of individuals contained more than 3 clones (Figure 4a). Surprisingly, 37% of polyclonal individuals were homogeneous for the ITD classification resulting in 90% of individuals harboring a single class of *FLT3*-ITD (Figure 4b), which may suggest a single ongoing mechanism of mutation even in the polyclonal samples. These observations implicate potentially distinct mechanisms in the etiology of the two classes of *FLT3*-ITD mutations or that they are enriched in particular genetic backgrounds.

As typical and atypical clones may have other distinct structural features beyond the exogenous nucleotides within the spacer segment, we sought to quantify these characteristics. We observed significantly longer duplications (Individuals: p = 0.011, Figure 4c; All clones: p = 0.007, Figure S1a) and spacers (Individuals: p <2.22e-16, Figure 4d; All clones: p <2.22e-16, Figure S1b) in atypical clones versus typical ones, which may point to the heterogeneity of these longer duplications in earlier studies (30,39,40).

We further characterized the exogenous region of ITD mutations. While the median length of the exogenous segment was 2 bp, we observed a clone with 51 exogenous nucleotides in our cohort (Figures 4e, and S1c). Upon further investigation, we found this segment originated from the intron 3’ of the *FLT3* exon 14. While we were able to clarify the origin of the exogenous segment of the ITD in this case, for the most part, smaller exogenous segments in other ITDs were highly promiscuous across the genome and it was challenging to resolve their origin. As such, we only characterized the nucleotide makeup of inserted segments to gain insight into biases of their nucleotide usage, and observed that the exogenous segments were significantly enriched for purines (p = 0.0112) and even more significantly for guanine nucleotides (G > A (p = 0.00559), G > C (p = 7.32e-5), G > T (p = 0.00940), and (p > 0.05) for all others, Figure S1d).

In our cohort, typical and atypical *FLT3*-ITD clones occurred with 77.6% and 22.4% frequencies, respectively. Observing divergence between the typical and atypical ITD features (Figures 4, and S1), we next investigated the impact of ITD structural characteristics on extending overall survival (OS) of 58 FLT3i-treated patients whom could be classified as *de novo* or relapsed AML (Table 1, see Material and Methods). We considered OS, defined as the time from diagnosis until death or last follow-up, as the clinical outcome. We used stratified Cox regression for the analysis to control for disease stage (*de novo* and relapsed) with sex, age, allogenic transplant, remission at initiation of TKI treatment, and FLT3i type as additional covariates.

We first investigated the impact of ITD structural characteristics on extending OS of FLT3i-treated patients. Previous studies documented an association of FLT3i treatment with improved OS in smaller cohorts but did not specifically evaluate the effect associated with the presence of *FLT3*-ITD mutations (18,19). Furthermore, there have been conflicting reports on whether *FLT3*-ITD duplication length is a significant prognostic factor in AML patients treated with standard chemotherapy. Several earlier studies (30,40) reported that longer duplication length significantly associated with inferior OS, while other studies did not find any significant link between the ITD length and OS of AML patients after standard chemotherapy (29,39). In our analysis, we found that length had no effect on the OS of FLT3i-treated AML patients (Cox PH: HR = 0.996, CI = 0.970 − 1.02, p = 0.756) whether these patients were newly diagnosed or were relapsed.

The lack of correlation between duplication length and survival incentivized the search for alternative sequence characteristics that could describe the heterogeneity of response to FLT3i treatment. We inquired whether the appearance of exogenous nucleotides within the ITD spacer segment impacted OS. When segregating the cohort into typical and atypical *FLT3*-ITDs, we found that carrying the typical ITD genotype reduced hazard by 76% (Cox PH: HR = 0.242, CI = 0.092 − 1.640, p = 0.0042, Figure 5a). We independently tested this observation in the TCGA cohort of newly diagnosed AML patients and observed a similar trend between typical and atypical *FLT3*-ITDs and OS in patients treated with standard chemotherapy alone without FLT3i, yet this result was marginally insignificant (Cox PH: HR = 0.425, CI = 0.170 − 1.06, p = 0.0660). However, the ITD classification significantly correlated with differential OS when the duplication length was taken into account as a covariate. While a significant correlation between the ITD classification and differential OS was observed for duplications longer than 18 bp (Cox PH: HR = 0.332, CI = 0.121 − 0.911, p = 0.0323), the most significant separation was seen for ITDs with 22 bp or longer duplication, where individuals with typical *FLT3*-ITDs had 76.3% reduced hazard compared to the patients with atypical ITDs (Cox PH: HR = 0.237, CI = 0.075 − 0.746, p = 0.0139, Figure 5b). Together, these data demonstrate that atypical ITD mutations lower the survival rate of AML patients.

Finally, to assess the role of spacer nucleotide composition on overall survival of FLT3i-treated patients, we examined differential OS in response to FLT3i as a function of the fraction of exogenous nucleotides in the ITD spacer. We found that increased fraction of the spacer sequences exogenous to the *FLT3* locus were positively correlated with lower OS rates (Cox PH: HR = 41.0, CI = 1.43 − 1180, p = 0.0302, Figure 5c). Repeating similar analysis for the TCGA AML cohort with duplication length as a covariate, we found that patients with a higher fraction of exogenous nucleotides exhibited 124 times increased hazard (Cox PH: HR = 124, CI = 1.99 − 7660, p = 0.0222, maximum separation obtained at 27 bp, Figure 5d). These data elucidate that the correlation of *FLT3*-ITD structure with prolonged survival in response to FLT3i is a multivariate relationship, potentially dependent on the length and composition of nucleotides residing within the spacer sequences. Taken together, these observations exhibit that the selectivity of response to FLT3i depends not only on the presence but also the nucleotide composition of ITD mutations, and AML patients with atypical FLT3 mutations on average have decreased survival in response to treatment with chemotherapy alone as well as chemotherapy with FLT3i.

## Discussion

In this study, we introduced two distinct classes of typical and atypical ITDs based on the mutation nucleotide composition, showed significant divergence between the characteristics of these two ITD classes, and developed an efficient and easy to use software (available at https://github.com/faryabib/HeatITup) to classify and annotate ITD mutations. Critically, our data demonstrate that responses to FLT3i treatment correlates with structural features and classification of *FLT3*-ITD mutations, and suggest that carrying atypical ITD mutations with nucleotides mainly exogenous to the *FLT3* sequences lowers the survival rate in AML patients from two independent cohorts with and without FLT3i treatment. We thus propose that the complexity of ITD mutations can be characterized and utilized to inform clinical decision-making and therapeutic algorithms for patients with AML in the era of molecular medicine. To this end, we have packaged HeatITup as an open source tool with the aim of assisting similar analysis in independent AML cohorts.

Although this study is a step toward implementing precision medicine for AML patients, further studies are warranted to fully understand the complexity of leukemogenesis in individual *FLT3*-ITD-positive patients, and further investigate the mechanism of this differential response. Our characterizations and similar observations underscore the importance of understanding interactions between *FLT3* and other recurrent mutations as well as physiological factors before attempting to predict clinical response to *FLT3*-based therapies.

While the crystal structure of *FLT3* suggests that ITDs negate the intrinsic auto-inhibitory activity of the juxtamembrane domain (JM) (8,41), it is unclear how highly heterogeneous characteristics of ITDs affect receptor function. One could hypothesize that more complex ITD mutations deviate further from the receptor structure. The long atypical ITDs with a significantly large fraction of exogenous nucleotides in their spacer segment might further disrupt the auto-inhibitory activity of the JM domain of the receptor, while small size duplications or less deviation from the endogenous locus may preserve some of the intrinsic auto-inhibitory function of the JM. Our correlative study suggests that the structure of the *FLT3*-ITD plays a prognostic role, but further studies, such as those in Smith *et al*. (42), are needed to clarify the mechanism of this effect and how it impacts leukemogenesis and drug response.

Recurrent ITD mutations are not unique to *FLT3* and have been reported in a related receptor KIT (8) and histone-lysine N-methyltransferase 2A (also known as mixed lineage leukemia) (KMT2A/MLL1) in acute lymphocytic leukemia (43). Also, ITD mutations are frequently observed in solid tumors such as BCOR in clear cell sarcoma of the kidney (44). Our results warrant investigation of the link between the complexity of recurrent ITD in genes beyond *FLT3* in AML and other cancers. Our algorithm, HeatITup, provides an accessible computational solution for accurate detection and classification of ITD that will enable these studies.

**Figure 1.**
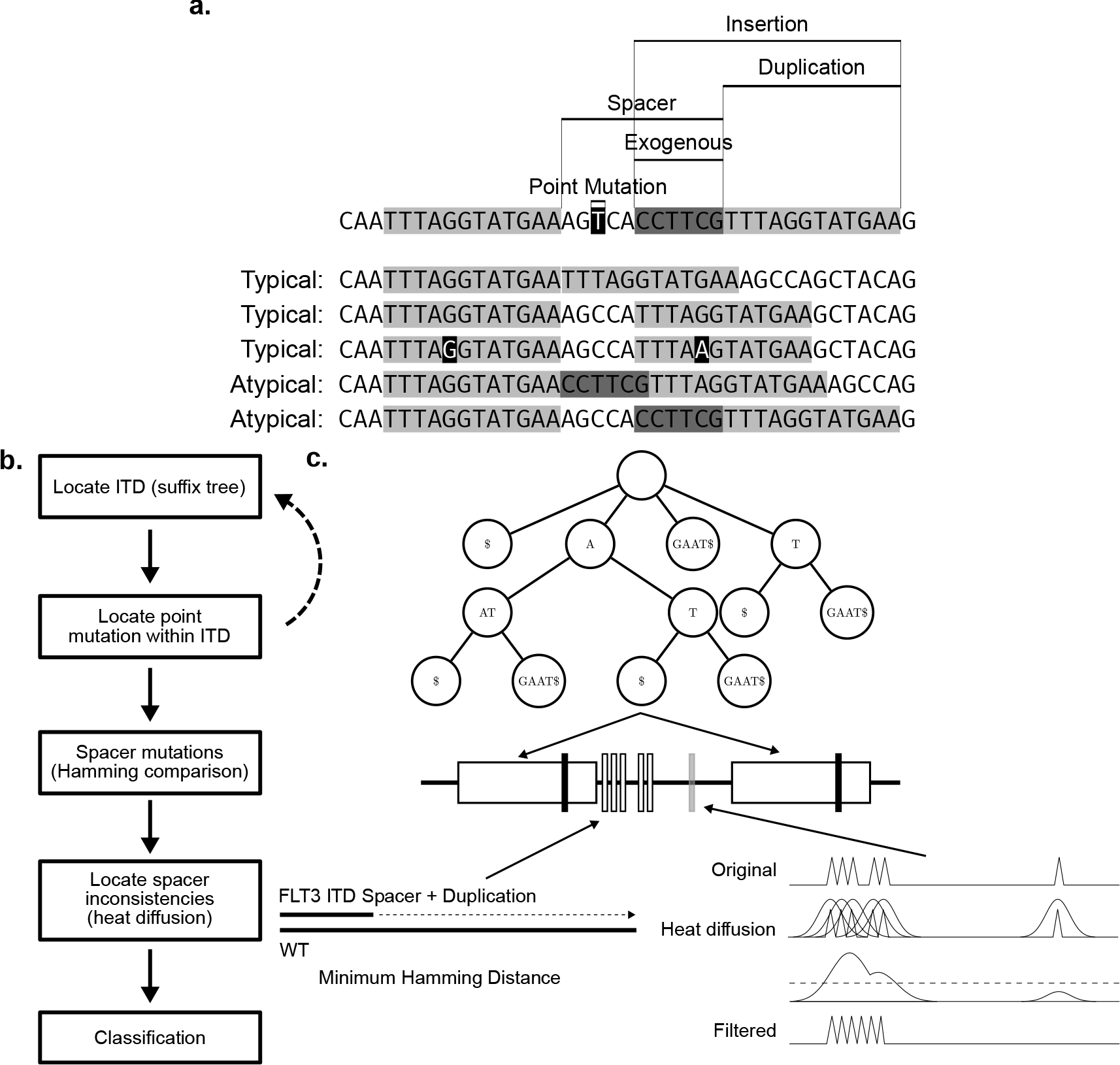
Schematic of ITD classification definition and HeatITup algorithm. a) Schematic of potential typical and atypical ITD mutations with marked duplication, spacer, and exogenous nucleotides. First sequence: structure of ITDs. Remaining sequences are examples of typical and atypical ITDs. Top to bottom: Duplication with no spacer; Duplication with spacer; Duplication with spacer and a point mutation; Duplication with entire spacer exogenous to exon; Duplication with part of spacer exogenous to exon. Light gray: duplicated, black: point mutation, dark gray: exogenous sequences. b) Outline of the HeatITup algorithm. c) Schematic of HeatITup modules and their function to analyze an ITD mutation. In this schematic, the *FLT3* locus ITD sequence (middle, black horizontal line) is composed of the repeated substring (duplication, in large white boxes), a point mutation within the duplication (vertical black bars), a point mutation within the spacer (vertical gray bar), and exogenous nucleotides (vertical white bars). The duplications with possible point mutations are found using a suffix tree (top). Mismatches in the spacer are found by scanning the duplicated segment with the spacer across the reference (bottom left) and exogenous nucleotides are found using the heat equation (bottom right).

**Figure 2.**
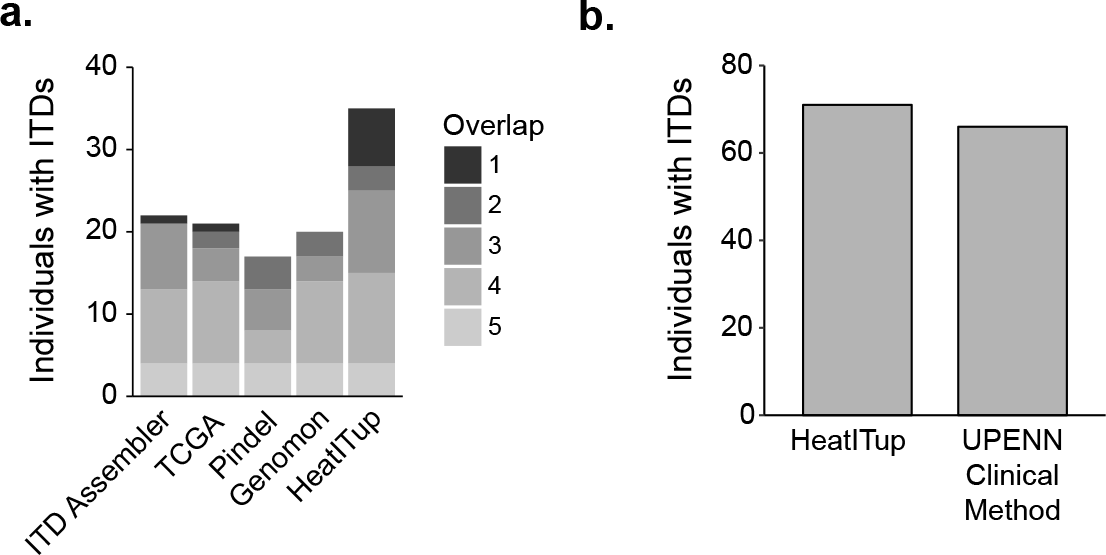
Comparison of HeatITup with commonly used ITD detection algorithms. Comparative analysis of HeatITup with a) four other methods on the TCGA AML cohort; or b) GATK/Pindel-based analysis workflow of CLIA-confirmed laboratory on the University of Pennsylvania cohort. In (a), fill color represents the number of algorithms that identified that *FLT3*-ITD-positive individual.

**Figure 3.**
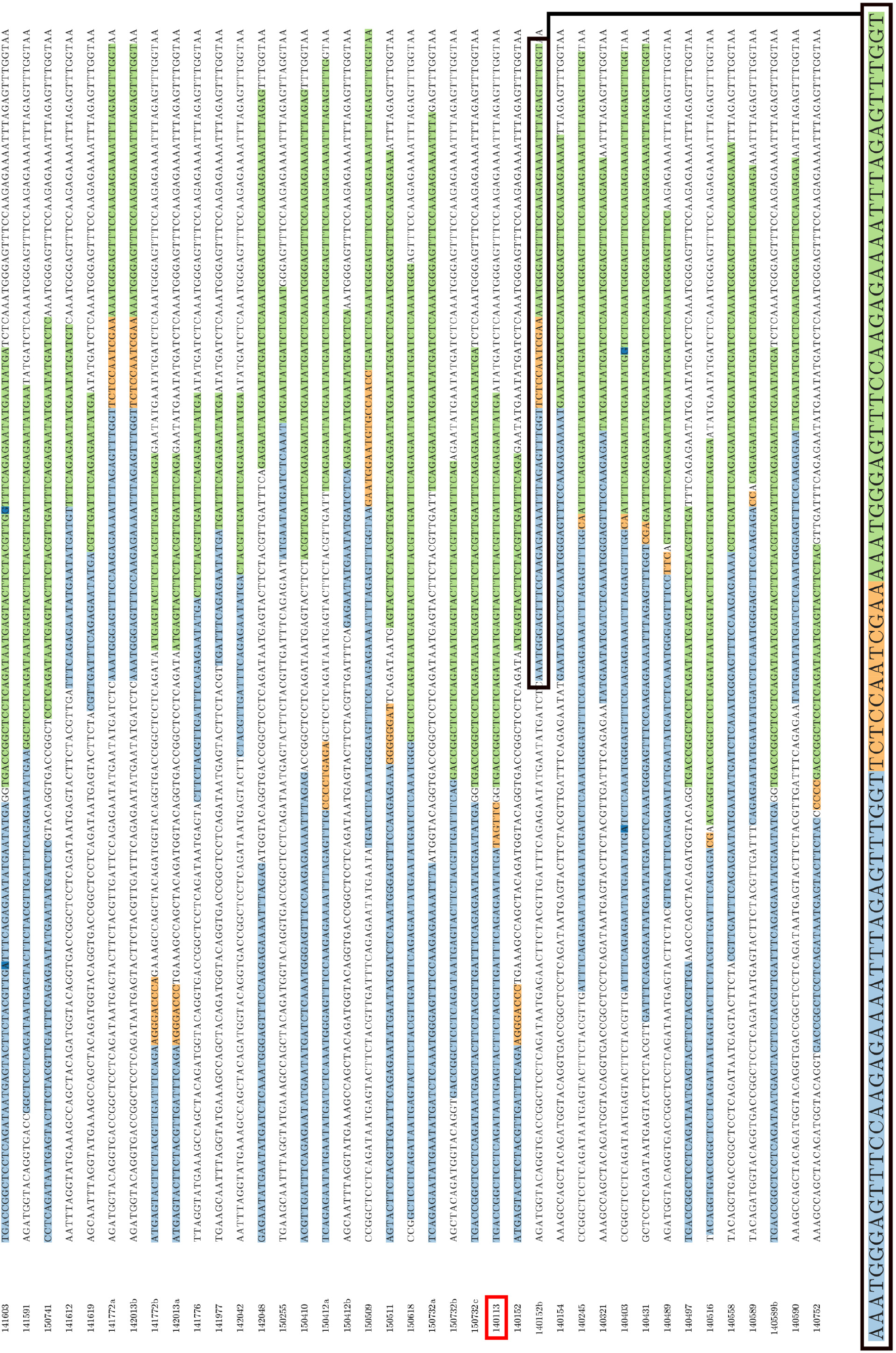
HeatITup-generated output automatically ITD annotates were uses for validation of *FLT3*-ITD classification. Light blue and green: the repeated sequence, dark blue: possible point mutations, orange: exogenous segments. Colors are customizable in the HeatITup rendered plot. Red box: CPDC140113, the manually misclassified sample. Letters indicate different clones in a sample. Black box: magnified ITD annotation.

**Figure 4.**
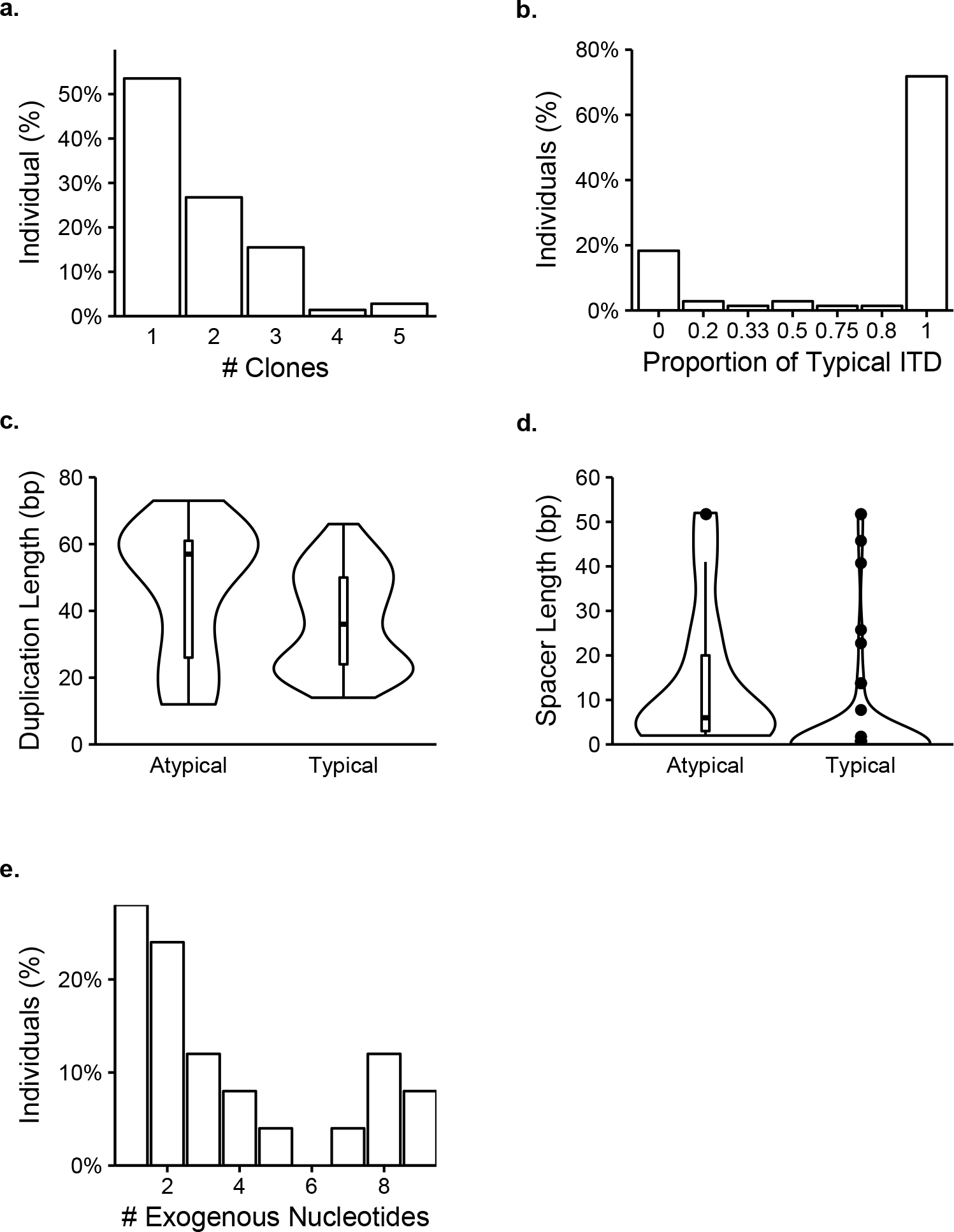
Characteristics of typical and atypical *FLT3*-ITDs nucleotide features. a)The percent of individuals per number of clones. b) The percent of individuals with different compositions of typical and atypical. The proportion of typical ITD represents (# typical clones / (# atypical clones + # typical clones)). c) Violin plots of duplicated segment lengths for typical and atypical individuals (Typical median: 36 bp, Atypical median: 57, Mann-Whitney U test: p = 0.0110). d) Violin plots of spacer segments for typical and atypical individuals (Typical median: 0, Atypical median: 6, Mann-Whitney U test: p < 2.22e-16). e) Distribution of the number of exogenous nucleotides per individual (median (mean): 2 (3.60) bp). n = 71 individuals.

**Figure 5.**
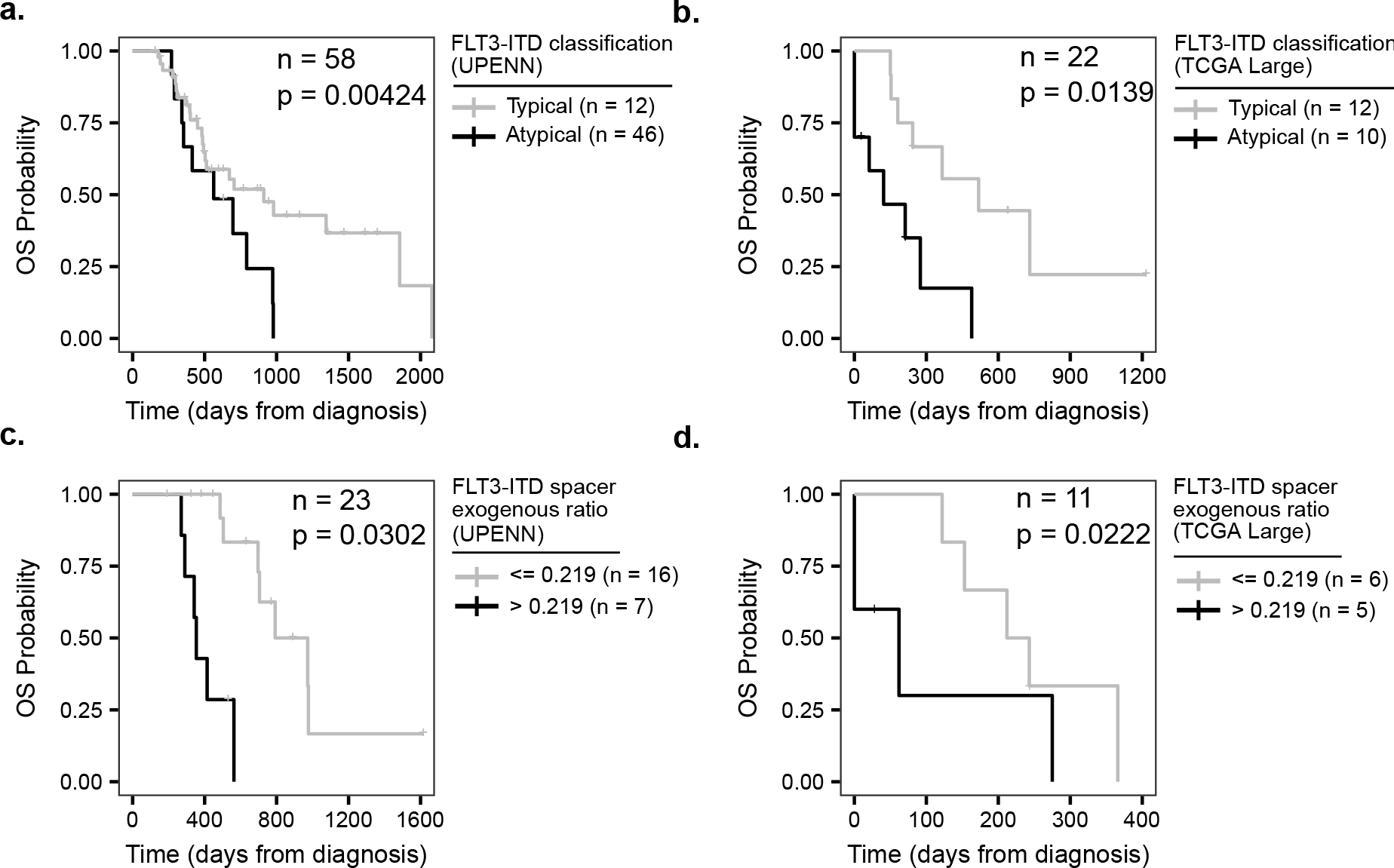
Correlation between ITD features and overall survival for the FLT3i-treated *FLT3*-ITD-positive AML patients. a) Typical (gray, median: 911 days) vs. atypical (black, median: 563 days) for all individuals in the University of Pennsylvania cohort (n = 58, Cox PH: HR = 0.242, CI = 0.092 − 1.640, p = 0.0042). b) Individuals with typical (gray, median: 518 days) vs. atypical (black, median: 122 days) clones in the TCGA AML cohort with duplication segment lengths > 22 bp (n = 14, Cox PH: HR = 0.237, CI = 0.075 − 0.746, p = 0.0139). c) Comparison of OS between individuals carrying ITDs with the exogenous ratio of > 0.219 (black, median: 354 days) with the ones with the exogenous ratio <= 0.219 (gray, median: 882 days). This ratio was determined by performing maximum separation analysis. The exogenous ratio was defined as (exogenous segment length / spacer segment length) for all spacer segment lengths (n = 23, Cox PH: HR = 41.0, CI = 1.43 − 1180, p = 0.0302). d) Individuals with duplication segment lengths > 27 bp with the exogenous ratio of > 0.219 (black, median: 62 days) vs. <= 0.219 (gray, median: 228 days) in the TCGA AML cohort as in (c) (n=11, Cox PH: HR = 124, CI = 1.99 − 7660, p = 0.0222). Cox regressions based on AML type (*de novo* or relapsed) with sex, age, allogenic transplant, remission at initiation of TKI treatment, and FLT3i type as additional covariates were used on the University of Pennsylvania cohort. In panels (c) and (d), exogenous ratio separations were only used for visualization and not Cox regression.

**Table 1.** Clinical characteristics of 58 *FLT3*-ITD positive acute myeloid leukemia patients treated with FLT3 inhibitors. ^A^ Of the 32 *de novo* patients, none had favorable cytogenetics, 29 had intermediate risk, 3 had adverse risk. ^B^ Patients were categorized into 4 major groups based on their FLT3i treatment. ^B^ Other FLT3i includes Quizartinib (3), LPX3397 (5), Midostaurin (1).

|  | n | % |
| --- | --- | --- |
| All Subjects | 58 | – |
| Sex |  |  |
| Male | 30 | 51.7 |
| Female | 28 | 48.3 |
| Age at Diagnosis |  |  |
| < 60 | 33 | 56.9 |
| ≥ 60 | 25 | 43.1 |
| Disease Status |  |  |
| De Novo<sup>A</sup> | 32 | 55.2 |
| Relapse | 26 | 44.8 |
| Induction Chemotherapy |  |  |
| Yes | 56 | 96.6 |
| No | 2 | 3.4 |
| FLT3i |  |  |
| Yes | 58 | 100 |
| No | 0 | 0.0 |
| FLT3 inhibitory drug |  |  |
| Sorafenib | 17 | 29.3 |
| Gilteritinib | 23 | 39.7 |
| Sorafenib and Gilteritinib | 9 | 15.5 |
| Others<sup>B</sup> | 9 | 15.5 |
| Allogeneic Transplant |  |  |
| Yes | 40 | 69.0 |
| No | 18 | 31.0 |
| Disease Status at initiation of TK |  |  |
| Remission | 16 | 27.6 |
| Active Disease | 42 | 72.4 |

